## Supplementary material for "LRRK2-mutant microglia and neuromelanin synergize to drive dopaminergic neurodegeneration in an iPSC-based Parkinson’s disease model": Supplental file

**Supplementary Table 1.** Description of antibodies used for Immunocytochemistry.

| Antibody | Host | Reactivity | Concentration | Source |
|---|---|---|---|---|
| CX3CR1 | Rabbit | Human | 1:200 | Abcam |
| FOXA2 | Goat | Human | 1:50 | R&D Systems |
| GFP | Chicken | Human, mouse | 1:250 | Aves Labs |
| IBA1 | Rabbit | Human, mouse | 1:200 | Wako |
| IBA1 | Mouse | Human | 1:200 | Santa Cruz |
| LMX1A | Rabbit | Human | 1:1000 | Millipore |
| MAP2 | Chicken | Human, mouse | 1:1000 | Abcam |
| TH | Rabbit | Human, rat | 1:500 | Sigma-Aldrich |
| TH | Mouse | Human, rat | 1:1000 | Merck Millipore |
| TMEM119 | Mouse | Human | 1:200 | Biolegend |
| TUJ1 | Mouse | Mammalian | 1:500 | Biolegen |

**Supplementary Table 2.** Description of antibodies employed for Flow cytometry.

| Antibody | Clone | Reactivity | Concentration | Source |
|---|---|---|---|---|
| CD11b-PE | M1/70 | Human, mouse | 1:50 | Biolegend® |
| CD14-Beads | - | Human | 1:20 | Miltenyi |
| CD163-APC | GHI/61.1 | Human | 1:50 | Miltenyi |

CD: Cluster of differentiation; PE: Picoeritrine; APC: Allophycocyanin.

**Supplementary Table 3.** Description of primers employed for gene expression analysis.

| Gene ID | Forward primer (5' - 3') | Reverse primer (5' - 3') |
|---|---|---|
| *C1Qa* | ATGGTGACCGAGGACTTGTG | GTCCTTGATGTTTCCTGGGC |
| *GAS6* | GTAGCTTCCACTGTTCCT | GCGCACTCGTCTATGTCTT |
| *GPR34* | GAAGACAATGAGAAGTCATACC | TGTTGCTGAGAAGTTTTGTG |
| *MerTK* | CTTCTCCATGGCCACAGGTT | ATACTGAAAAGGTGGGGCGG |
| *P2RY12* | CTAAGATTCTCTGTTGTCATCTG | ACAGAGTGCTCTCTTTCACATAG |
| *PROS1* | AAGAAGCCAGGGAGGTCTTTG | ACGTGCAGCAGTGAATAACC |
| *IL-6* | AATTCGGTACATCCTCGACGG | GGTTGTTTTCTGCCAGTGCC |
| *C3* | AAAAGGGGCGCAACAAGTTC | GATGCCTTCCGGGTTCTCAA |
| *IFNγ* | CATTACCTGAAGGCCAAGGA | CTGACTATGGTCCAGGCACA |
| *β-Actin* | AGGCCAACCGCGAGAAG | ACAGCCTGGATAGCAACGTACA |

**Supplemental Table 4.** Description of the antibodies used for Western Blot.

| Antibody | Host | Reactivity | Concentration | Source |
|---|---|---|---|---|
| PCNA | Mouse | Human | 1:3000 | Sigma-Aldrich |
| PSD-95 | Mouse | Human | 1:1000 | Synaptic Systems |
| β-Actin | Mouse | Human | 1:2000 | Affinity Biosciences |

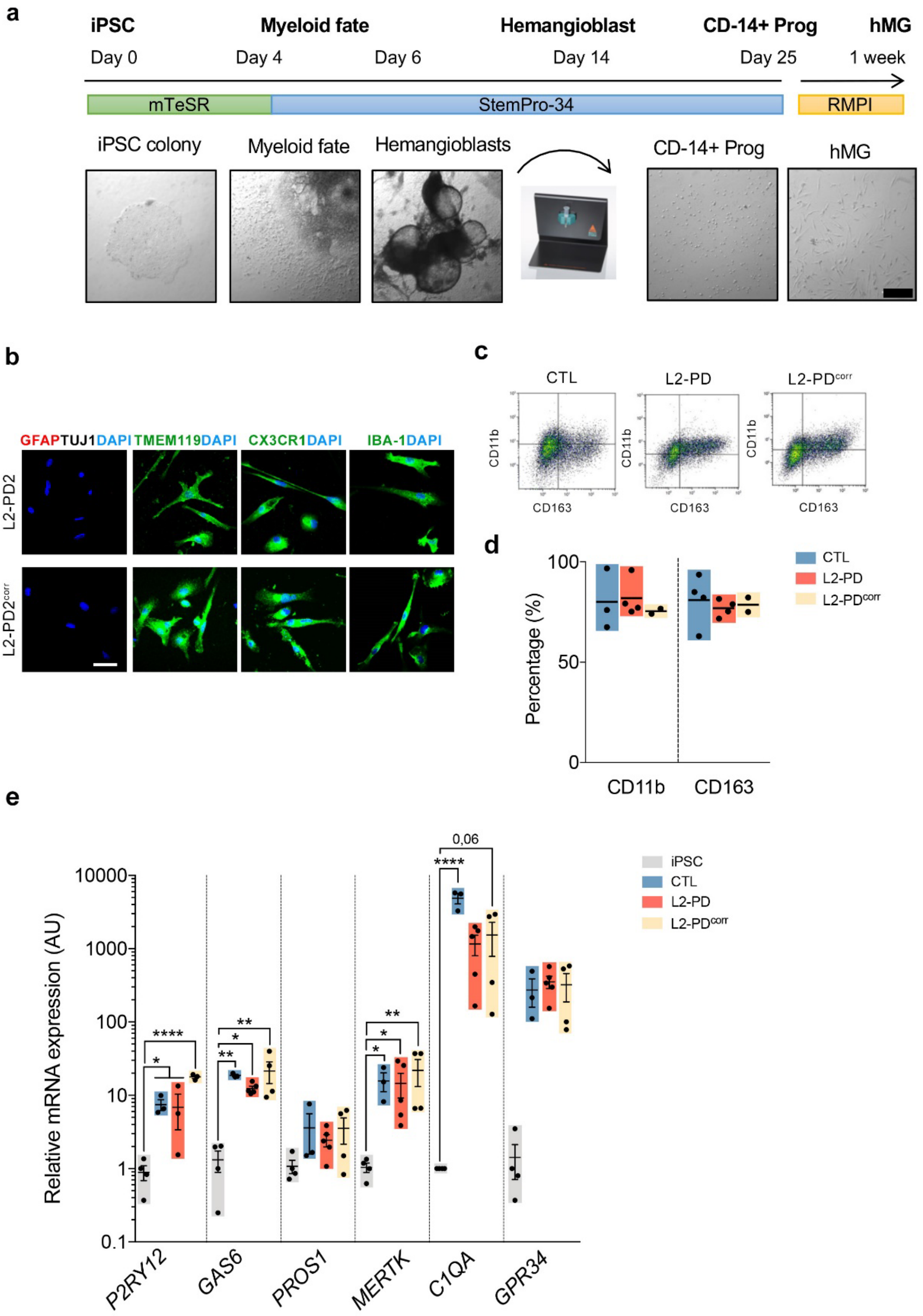


**Supplementary Figure 1. Generation and characterization of hMG derived cells. a** Schematic representation of the differentiation protocol to generate iPSC-derived hMG with representative bright field pictures of the process (Scale bar=100 μm). **b** Representative immunocytochemistry (ICC) images of iPSC-derived hMG after 7 days in culture from L2-PD2 (SP13) and L2-PD2$^{corr}$ (SP13wt/wt) iPSC lines staining positive for IBA-1, CX3CR1 or TMEM-119 (green) and negative for astrocytic (GFAP) or neuronal (TUJ1) markers. Nuclei are counterstained with DAPI (blue). Scale bar=30 μm. **c-d** Representative flow cytometry plots

for CD11b and CD163 microglial surface markers from CTL (SP09), L2-PD (L2-PD2: SP13) and L2-PD$^{corr}$ (L2-PD2$^{corr}$: SP13wt/wt) and its corresponding quantification. Individual data plotted, along with mean ± SEM. N=3 for CTL, N=4 for L2-PD, and N=2 L2-PD$^{corr}$. Intact cells were gated in a forward and side scatter (FSC/SSC) plot to exclude small debris. Gating of live cells was done using the viability dye PI. **e** Relative mRNA expression of human specific microglia genes comparing iPSCs with hMG from CTL (SP09), L2-PD (L2-PD1: SP12; L2-PD2: SP13) and L2-PD$^{corr}$ (L2-PD1$^{corr}$: SP12wt/wt; L2-PD2$^{corr}$: SP13wt/wt). Individual data plotted, along with mean ± SEM. One-way ANOVA with Uncorrected Fisher LSD test for all comparison, except Kruskal-Wallis non parametric test with Uncorrected Dunn's test for *PROS1* and *MERKT* comparisons. N=4 for iPSC, N=3 for CTL, N=5 for L2-PD, and N=4 for L2-PD$^{corr}$.
*p<0.05, **p<0.01, ***p<0.001, ***p<0.0001; p-value is specified for values between 0,05 and 0,1. p-values over 0.1 (non-significant) are not shown.

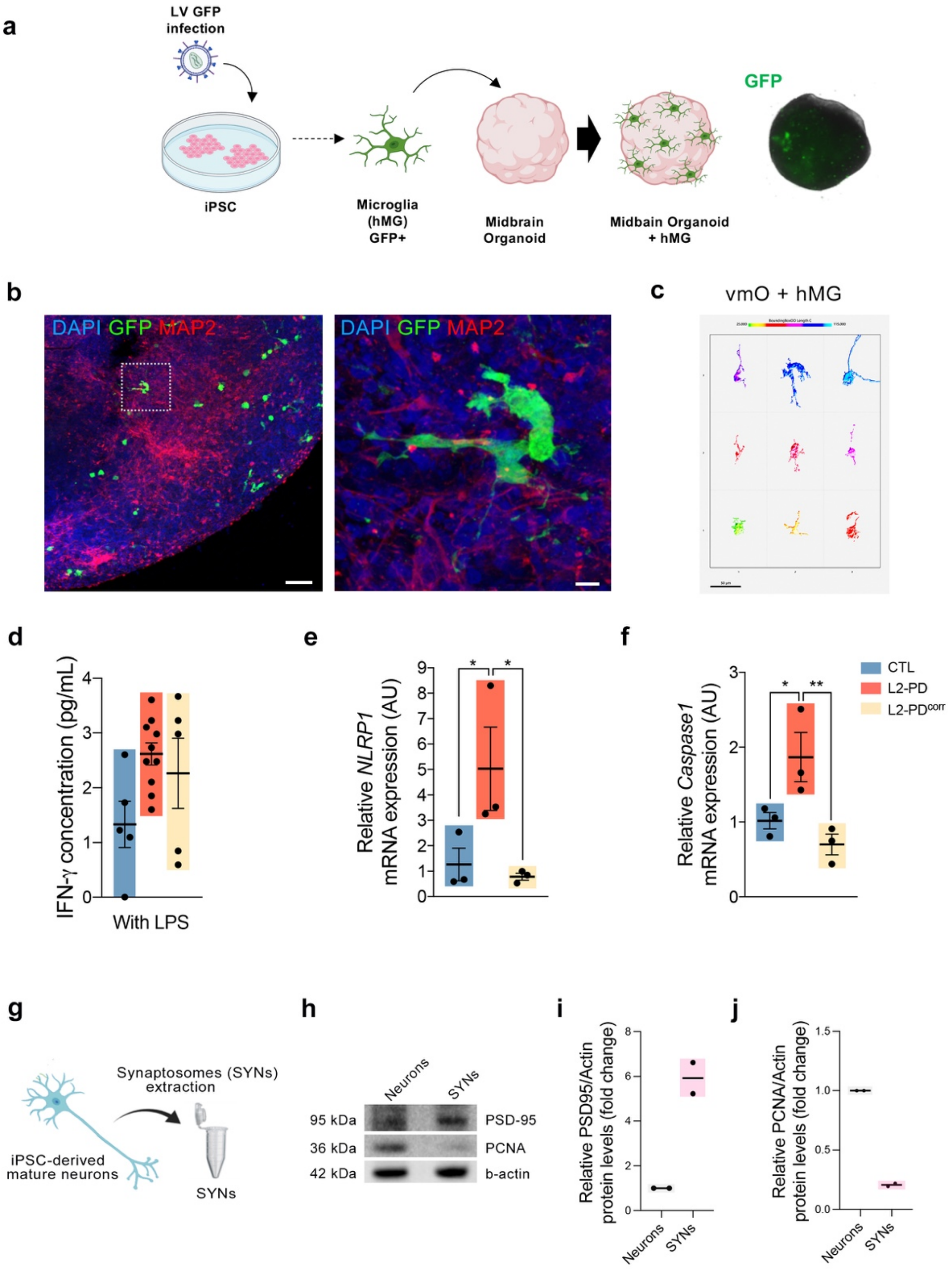


**Supplementary Figure 2. 3D co-culture set up, hMG morphological analyses and functional validation. a** Schematic representation of the 3D co-culture procedure. **b** Representative ICC images of iPSC-derived CTL hMG (SP09, green) inside vmO from a CTL line (SP11). Mature neurons are stained with MAP2 (red) (Scale bars=100 and 10 µm). **c** Representative IMARIS reconstructions from IN TOTO ICC of CTL hMG (SP09) showing ramified morphologies (Scale bar=50 µm). **d** Cytokine profile for IFN-y after 24 hours of LPS stimulation in CTL (SP09), L2-PD (L2-PD1: SP12; L2-PD2: SP13) and L2-PD2$^{corr}$ (SP13wt/wt) hMG. Individual data plotted, along with mean ± SEM. N=3 of independent experiments, each experiment containing two technical duplicates. **e-f** Relative mRNA expression of inflammasome-related genes after 24h of LPS stimulation of CTL (SP09), L2-PD (L2-PD1: SP12) and L2-PD$^{corr}$ (L2-PD1$^{corr}$: SP12wt/wt) hMG. Individual data plotted, along with mean ±

SEM. One-way ANOVA with Uncorrected Fisher LSD test. N=3 of independent experiments. **g** Schematic representation of Synaptosome extraction from iPSC-derived mature neurons. **h** Representative Western blot images for Postsynaptic Density protein (PSD)-95, Proliferating Cell Nuclear Antigen (PCNA), and β-Actin protein bands from iPSC-derived Mature Neurons (Neurons) and extracted SYNs. **i-j** Quantification of PSD-95 and PCNA relative to β-Actin, represented as Fold change to Neurons. Individual data plotted, along with mean. N=2 experiments. *p<0.05, **p<0.01. p-values over 0.1 (non-significant) are not shown.

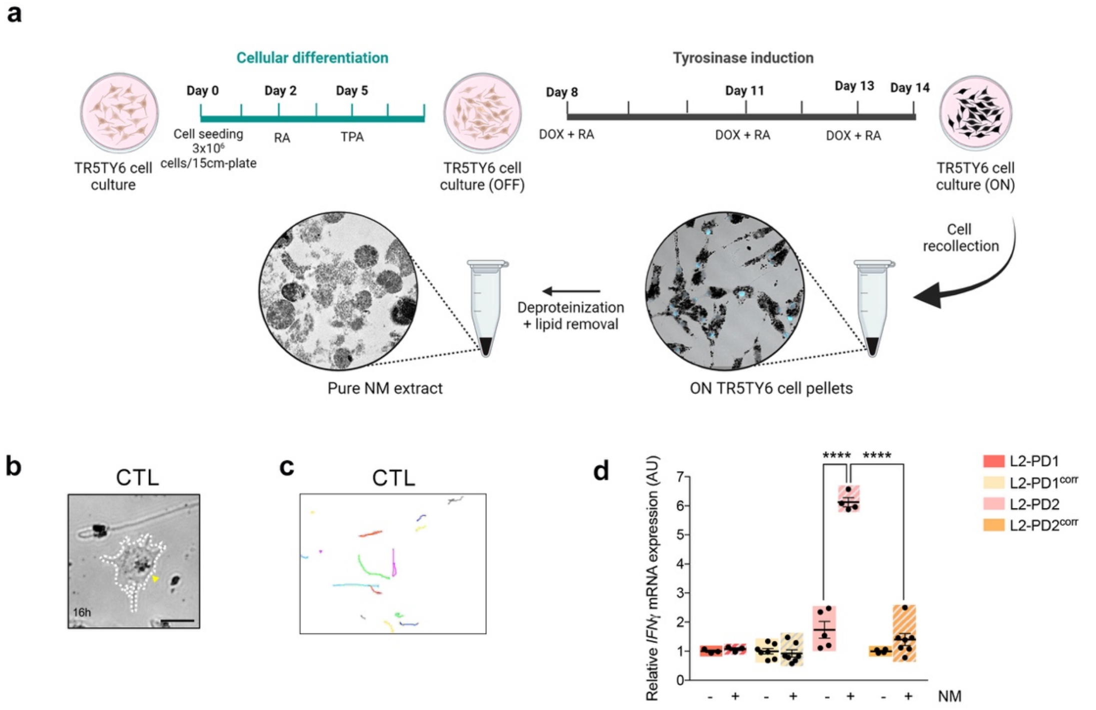


**Supplementary Figure 3. NM purification from cultured TR5TY6 neuroblastoma cells. a.** TR5TY6 neuroblastoma cells were differentiated for 6 days using retinoic acid (RA) and 12-O-tetradecanoylphorbol-13-acetate (TPA) and induced for human tyrosinase expression with doxycycline. Six days after induction, cells were collected and pure NM extracts were obtained after deproteinization and lipid removal. **b** Representative Bright Field images of CTL (SP09) hMG phagocyting NM particles for 16 hours (yellow arrow-heads for phagocyted particles; Scale bar=25 µm). **c** Spontaneous migration paths of CTL (SP09) hMG were tracked for 16h. The location of each cell was determined every 2 minutes and connected to depict its migration route. **d** Relative mRNA expression of IFN-y in L2-PD1 (SP12), L2-PD1$^{corr}$ (SP12wt/wt), L2-PD2 (SP13), and L2-PD2$^{corr}$ (SP13wt/wt) hMG at basal conditions or under NM stimulation for 24 hours. Individual data plotted, along with mean ± SEM. One-way ANOVA with Tukey multiple comparison test. N=3 of independent experiments, each experiment containing two technical duplicates. ****p<0.0001. p-values over 0.1 (non-significant) are not shown.

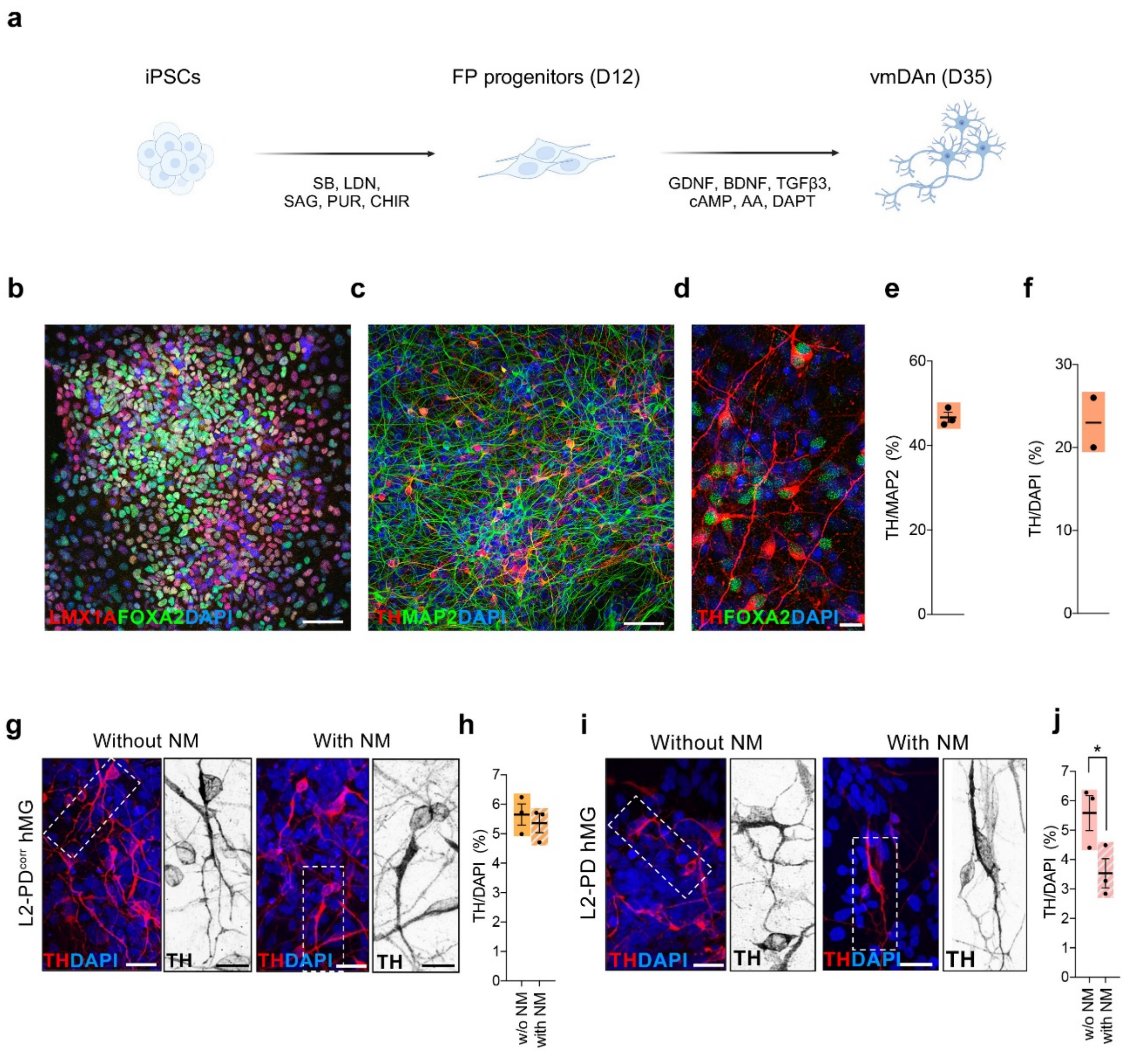


**Supplementary Figure 4. vmDAn characterization and degeneration upon culture with L2-PD hMG and NM. a** Schematic representation of the differentiation protocol employed to generate iPSC-derived vmDAn. **b** Representative immunocytochemistry (ICC) images of D12 CTL floor plate (FP) progenitors (SP11) staining positive for LMX1A and FOXA2. Nuclei are counterstained with DAPI (blue). Scale bar=65 µm. **c-d** Representative immunocytochemistry (ICC) images of D35 vmDAn (SP11) staining positive for TH and MAP2 (**c**) and TH and FOXA2 (d). Scale bars=50 and 13 µm. Nuclei are counterstained with DAPI (blue). **e-f** Quantification of percentage of TH/MAP2 (**e**) and TH/DAPI (**f**) in a monoculture of D35 CTL vmDAn (SP11). Individual data plotted, along with mean ± SEM. N=3 independent experiments (**e**), N=2 independent experiments (**f**). **g** Representative images of CTL TH+ vmDAn (SP11) upon culture with L2-PD2$^{corr}$ hMG (SP13wt/wt), without or with NM (Scale bar=25 µm; 15 µm). **h** Quantification of percentage of TH+ population over the total number of DAPI+ cells in culture with L2-PD2$^{corr}$ hMG (SP13wt/wt), without or with NM. Individual data plotted, along with mean ± SEM. Paired t test. N=3 independent experiments. **i** Representative images of CTL TH+ vmDAn (SP11) upon culture with L2-PD2 hMG (SP13), without or with NM (Scale bar=25 µm; 15 µm). **j** Quantification of percentage of TH+ population over the total number of DAPI+ cells in culture with L2-PD2 hMG (SP13), without or with NM. Individual data plotted, along with mean ± SEM. Paired t-test. N=3 independent experiments. *$p<0.05$. p-values over 0.1 (non-significant) are not shown.

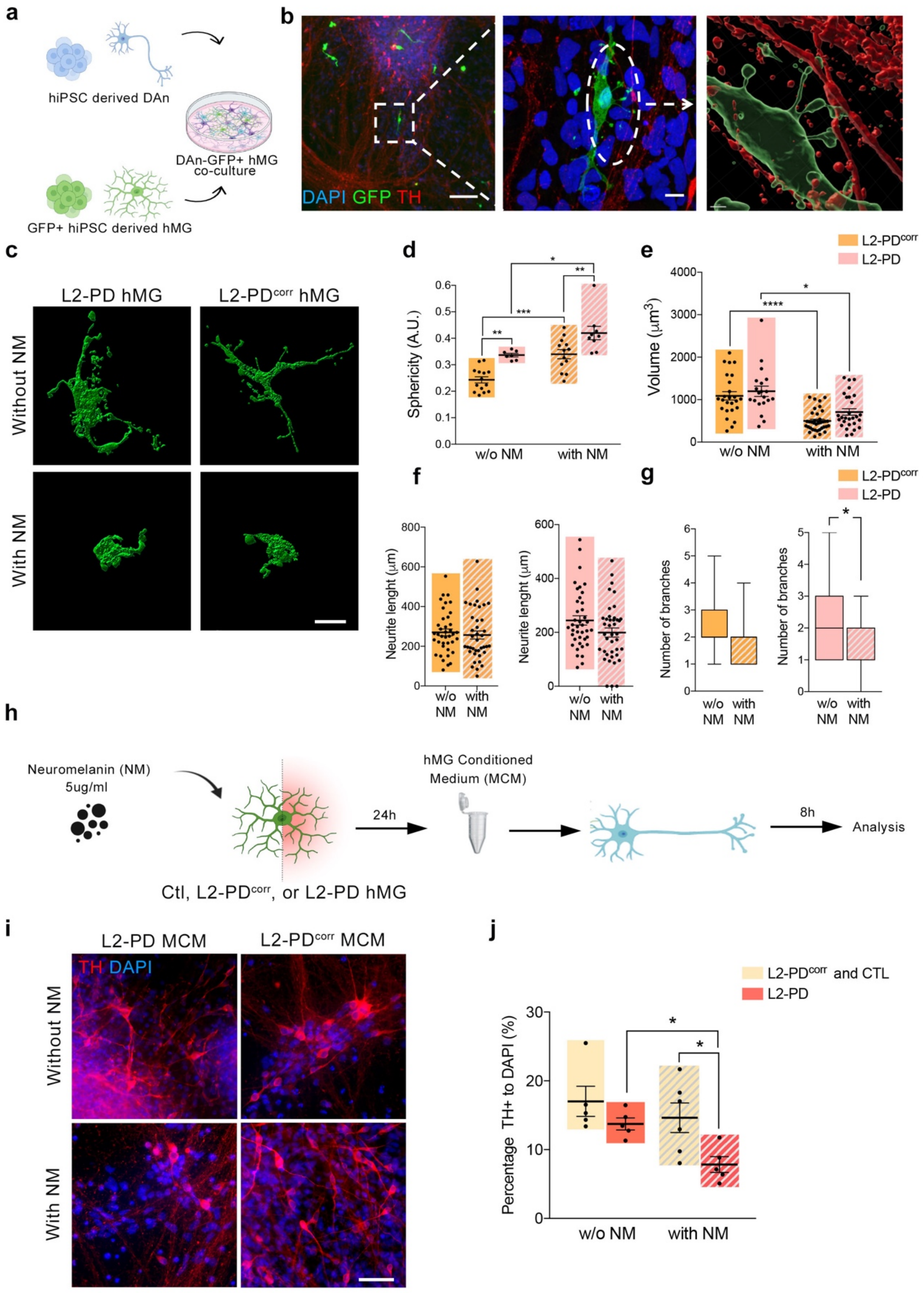


**Supplementary Figure 5. Set up of iPSC-derived 2D Neuron/Microglia co-culture system. a** Schematic representation of generation of a co-culture between vmDAns and GFP+ hMG. **b** Representative images of hMG (GFP+, green) in contact with CTL (SP11) TH+ vmDAns (red), with corresponding IMARIS reconstructions. Scale bar=70 μm; 7 μm; 2 μm. **c** IMARIS reconstructions from 2D Neuron/Microglia co-cultures of L2-PD (L2-PD2: SP13) and

L2-PD$^{corr}$ (L2-PD2$^{corr}$: SP13wt/wt) hMG showing ramified morphologies under basal condition and amoeboid shapes after NM stimulation (Scale bar=20 µm). **d-e** Quantification of microglial sphericity (**d**; AU) and volume (**e**; µm3) with IMARIS, comparing L2-PD2$^{corr}$ (L2-PD2$^{corr}$: SP13wt/wt) and L2-PD (L2-PD2: SP13) hMG, without or with NM. Individual data plotted, along with mean ± SEM. One-way ANOVA with Tukey multiple comparison test for (**d**); Kruskal-Wallis non-parametric test with Dunn multiple comparison test for (**e**). In (**d**) N=14 cells in L2-PD$^{corr}$, N=13 cells in the L2-PD$^{corr}$+NM, N=7 cells in L2-PD, N=9 in L2-PD+NM. The four conditions were measured from two independent experiments. In (**e**) N=25 cells in L2-PD$^{corr}$, N=36 cells in the L2-PD$^{corr}$+NM, N=20 cells in L2-PD, N=28 in L2-PD+NM. The four conditions were measured from two independent experiments. **f** Quantification of CTL vmDAns (SP11) neurite length (µm) upon co-culture with L2-PD$^{corr}$ (L2-PD2$^{corr}$: SP13wt/wt) or L2-PD hMG (L2-PD2: SP13), under basal condition or after stimulation with NM. Individual data plotted, along with mean ± SEM. Mann-Whitney test. In (**f**) N=39 cells in L2-PD$^{corr}$, N=39 cells in the L2-PD$^{corr}$+NM, N=40 cells in L2-PD, N=40 in L2-PD+NM. The four conditions were measured from two independent experiments. **g** Quantification of the number of terminals in CTL vmDAns (SP11) upon co-culture with L2-PD$^{corr}$ (L2-PD2$^{corr}$: SP13wt/wt) or L2-PD (L2-PD2: SP13) hMG, under basal condition or after stimulation with NM. Box and whiskers plot, depicting minimum and maximum value. Mann-Whitney test for L2-PD$^{corr}$ hMG; unpaired t-test for L2-PD hMG. In (**g**) N=40 cells in L2-PD$^{corr}$, N=39 cells in the L2-PD$^{corr}$+NM, N=39 cells in L2-PD, N=39 in L2-PD+NM. The four conditions were measured from two independent experiments. **h** Schematic representation of DAn treated for 8 hours with MCM exposed to NM (5ug/ml). **i** Representative ICC images of TH+ CTL vmDAns (SP11, Red) treated with either MCM from L2-PD (L2-PD2: SP13) and L2-PD2$^{corr}$ (L2-PD2$^{corr}$: SP13wt/wt) non stimulated or exposed to NM for 8 hours. Scale bar=50 µm. **j** Quantification of percentage of TH+ population over DAPI upon treatment with MCM from control or L2-PD (L2-PD1:SP12 and L2-PD2: SP13) hMG, non-stimulated or exposed to NM for 8 hours. CTL (SP09) and L2-PD$^{corr}$ (SP13wt/wt) hMG have been pooled together and considered as control group. Individual data plotted, along with mean ± SEM. Kruskal-Wallis non-parametric test with Uncorrected Dunn's test. For L2-PD$^{corr}$ and CTL lines at basal conditions N=5 measurements coming from two independent experiments. For L2-PD at basal conditions and with NM N=5 measurements coming from two independent experiments. For L2-PD$^{corr}$ and CTL with NM N=6 measurements coming from two independent experiments.

*p<0.05, **p<0.01, ***p<0.001, ***p<0.0001. p-values over 0.1 (non-significant) are not shown.

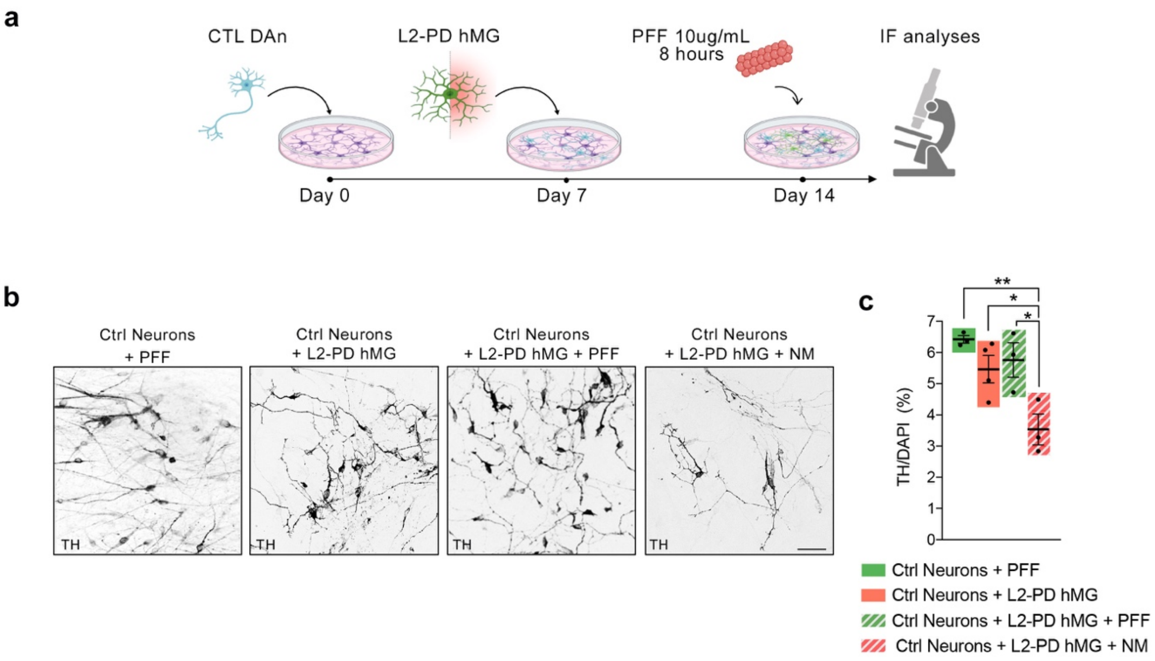


**Supplementary Figure 6. L2-PD hMG exposed to α-synuclein pre-formed fibrils (PFF) does not trigger DAn degeneration in an iPSC-derived 2D neuron/hMG co-culture system. a** Schematic representation of the co-culture system and PFF stimulation (10ug/ml). **b** Representative images of TH+ (black) CTRL (SP11) vmDAns treated with PFF, or in co-culture with L2-PD (L2-PD2: SP13) hMG treated with either PFF or NM (5ug/ml) (Scale bar=50 µm). **c** Quantification of percentage of TH+ population over the total number of DAPI+ cells. Individual data plotted, along with mean ± SEM. One-way ANOVA with Tukey multiple comparison test. N=4 for Ctrl Neurons + L2-PD hMG and N=3 for the other three conditions. *p<0.05, **p<0.01. p-values over 0.1 (non-significant) are not shown.
